# Assessing the impact of roadkill on the persistence of wildlife populations: a case study on the giant anteater

**DOI:** 10.1101/2022.01.18.476626

**Authors:** Fernando Ascensão, Arnaud L.J. Desbiez

**Author notes:** **Corresponding author:** Fernando Ascensão. Contact; cE3c – Centre for Ecology, Evolution and Environmental Changes, Faculdade de Ciências da Universidade de Lisboa, Edifício C2, 5° Piso, Sala 2.5.46 Campo Grande 1749-016 Lisboa, Portugal; (Ext. 22577).

## Abstract

Human activity is depleting biodiversity, and road networks are directly contributing to this trend due to roadkill. Nevertheless, few studies empirically estimated the impact of roadkill on wildlife populations. We integrated information on roadkill rates, population abundance, and animal movement to estimate the survival rates and the proportion of the population likely to be extirpated due to roadkill, using giant anteater (*Myrmecophaga tridactyla*) as model species. We then assessed the consequent implications of roadkill on population persistence using population viability analysis (PVA). The yearly survival rate of resident anteaters inhabiting road vicinity areas (0.78; CI:0.62-0.97) was considerably lower than for those living far from roads (0.95; CI:0.86-1.00). The real number of anteaters being road-killed is considerably higher than the one recorded in previous studies (by a factor of 2.4), with ca. 20% of the population inhabiting road vicinity areas being road-killed every year. According to PVA results, roadkill can greatly affect the persistence of the giant anteater populations by reducing the growth rate down to null or negative values. This study confirms that roads have significant impacts on local population persistence. Such impacts are likely to be common to other large mammals, calling for effective mitigation to reduce roadkill rates.

## Introduction

Billions of animals of various species are killed every year in wildlife vehicle collisions (WVC), with impressive estimates reaching 2.2 million mammals on Brazilian roads (González-Suárez et al., 2018) and 29 million mammals in Europe (Grilo et al., 2020). Virtually all species inhabiting road vicinity areas can be impacted by WVC, and such pervasive impact is reflected in the growing number of roadkill studies across the globe (Barrientos et al., 2021; Schwartz et al., 2020). However, to date there is still scarce information regarding the effects of roadkill on wildlife population persistence (Barrientos et al., 2021), and the key question remains: can the added mortality from WVC represent a threat that may lead to local population extinctions? This information is particularly relevant for larger mammals, known to be highly vulnerable to roadkill impacts (Fahrig and Rytwinski, 2009; Rytwinski and Fahrig, 2012).

To correctly assess the impact of WVC on wildlife population persistence, one requires a detailed knowledge of local species abundance and on roadkill rates (Visintin et al., 2016). However, compiling such information for the same species has seldom been achieved (e.g., D’Amico et al., 2015; Santos et al., 2016), precluding our understanding on the roadkill impact on wildlife species (Barrientos et al., 2021). Here, we used the giant anteater (*Myrmecophaga tridactyla*) as a model species to assess the real impact of WVC on local populations of a large-sized mammal. This species has a body mass of 33 kg, low recruitment, with about one pup per year (Gaudin et al., 2018), and low densities of < 1 ind./km^2^ (Bertassoni et al., 2021), traits that make them particularly vulnerable to additional non-natural mortality (Fahrig and Rytwinski, 2009; Rytwinski and Fahrig, 2012). Despite its wide distribution throughout Central and South America, this species is listed as “Vulnerable to Extinction”, and one of the recognized threats is being recurrently involved in WVC throughout its distribution (Miranda et al., 2014), namely in the Brazilian state of Mato Grosso do Sul (Ascensão et al., 2021, 2017).

We integrated and updated different sources of empirical and bibliographic information on giant anteaters, namely roadkill rates, local population, and telemetry data. We aimed to obtain credible estimates on *i*) the probability of survival of giant anteaters inhabiting road vicinity areas in comparison to those living far from roads; and on *ii*) the proportion of animals that are annually road-killed on main roads. We then used this information on population viability analysis to *iii*) assess the consequences of roadkill for anteater population persistence in road vicinity areas. We hypothesized that those anteaters living near paved roads have lower survival rates, given the high roadkill rates, and consequently a large proportion of anteaters was expected to be road-killed. Also, we expected to estimate a high proportion of individuals that are annually road-killed, overall having a significant impact on the persistence of anteaters by lowering their population growth rate, and consequently their resilience to overcome stochastic catastrophes.

## Methods

### Study area and datasets

The empirical data analyzed here was collected by the ‘Anteater & Highways Project’ (www.giantanteater.org), which has been running for the past five years in the state of Mato Grosso do Sul (MS), in the Cerrado biome (savannah) of Brazil. The project has focused on four highways (all two-lane roads, total 1160 km) and adjacent areas (Fig.1A). Traffic volumes (vehicles per day) of studied roads range approx. between 600 in MS040 (Fig.1D; *pers. obs*) and 4300 in BR267 (Fig.1E) (DNIT, 2020). In BR262, the traffic volume is ca. 3350 in the west section (Fig. 1B) and 3850 in the east section (Fig.1C) (DNIT, 2020) (see for further details on study area in Ascensão et al. 2021 and Noonan et al. 2021).

**Figure 1.**
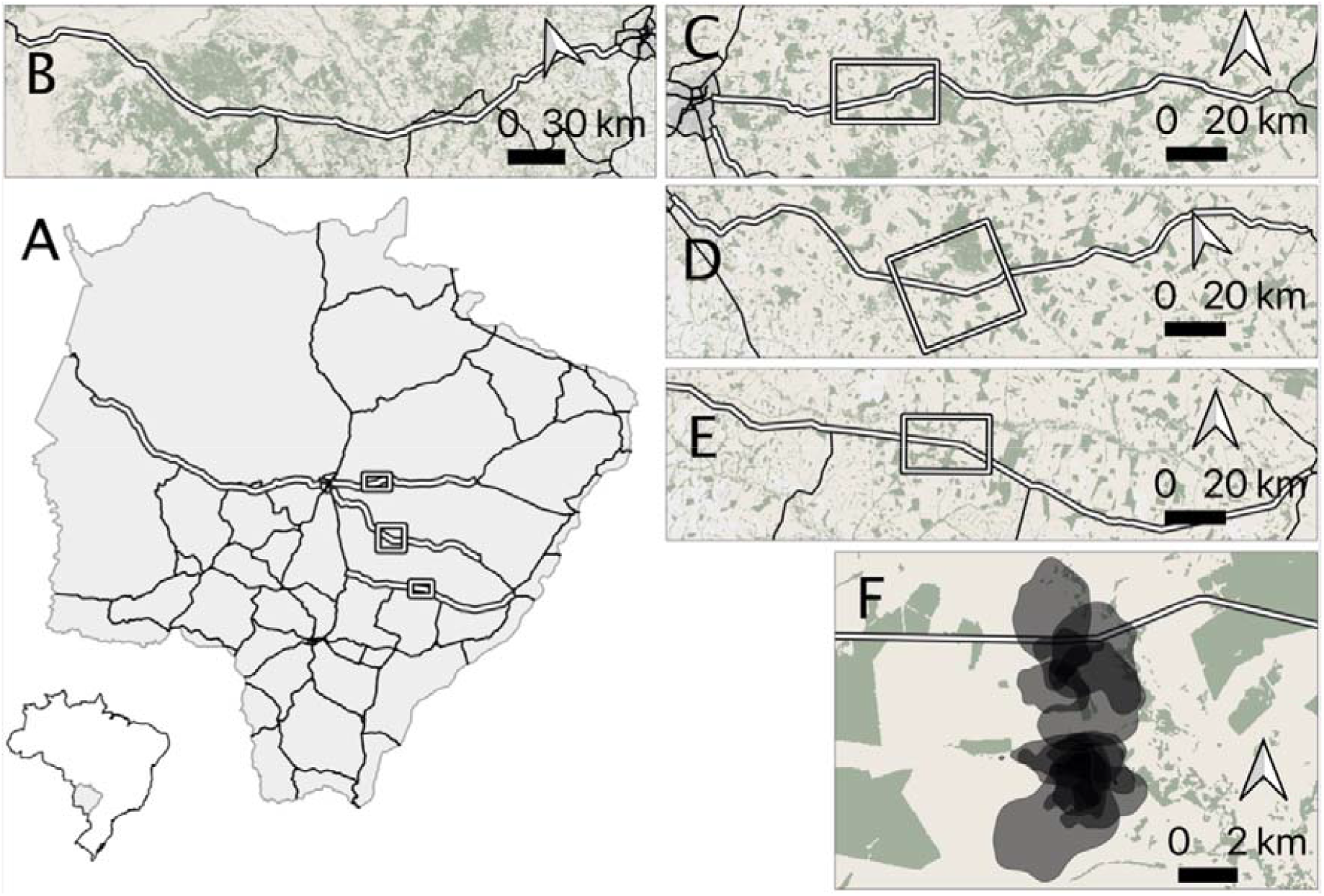
Location of the Brazilian state Mato Grosso do Sul and the main road network (A). Arrows indicate the different study areas where roadkill (B-E) and movement (C-E) has been collected within the research project ‘Anteater and Highways’ (www.giantanteater.org). Green areas correspond to native forest patches and yellow areas are mostly pasture areas. In B, C, D and E, the white lines indicate the roads BR-262 (west), BR-262 (east), MS-040, and BR-267, respectively. In F is shown the home ranges of 12 anteaters tracked in MS-040 (D).

### Roadkill rates

Roadkill data has been systematically collected throughout the four main paved roads (Fig. 1B-E) (Ascensão et al., 2021, 2019b, 2017). Here, we used the dataset from Ascensão et al. (2021). Briefly, roadkill surveys were carried out by car (40-60 km/h), on the four roads, searching for road-killed animals on both lanes and shoulders, for three years, on a fortnightly basis. A total of 420 survey events were performed, totaling 84,673 km of survey effort. During this study, 608 road-killed giant anteaters were recorded, which represented a mean annual roadkill rate of ∼0.19 ind./km/year (raw values) across surveyed roads. In parallel with roadkill surveys, carcass persistence experiments were also carried out to estimate the roadkill rate more accurately by accounting for carcass persistence and detectability bias. Experiments were performed on three of the surveyed roads, one with low traffic volume (MS040) and two with higher traffic volume (BR267 and BR262-east), with daily surveys for 15-30 days (mean 25.9 days). The carcasses persistence information from 21 anteaters plus of 96 individuals pertaining to species having a similar weight as giant anteaters (67 *Cerdocyon thous*, 21 *Hydrochoerus hydrochaeris*, 7 *Tapirus terrestris* and 1 *Mazama gouazoubira*) were used to correct the roadkill rates [see Ascensão et al. (2021) for further details on carcass persistence data collection].

### Animal movement information

Movement data of giant anteaters has been collected for the past five years (2017-2021) in the vicinity of three of the paved roads (Fig. 1C-E) (Noonan et al., 2021); currently totaling 46 adult wild giant anteaters (25 females and 21 males). In this dataset, four anteaters (ca. 9%) are known to have dispersed during their tracking period, of which two died while dispersing (one road-killed on a dirt road and one from unknown cause), and the other two settled in a new territory after dispersing. Other seven (all resident) individuals were also road-killed during the tracking period, one of which inhabited a territory far from paved roads but was hit on a dirt road. Overall, of 46 animals monitored, at least eight (17%) died due to vehicle collisions, with two collisions occurring on dirt roads (far from paved roads), and six on paved roads. The anteaters were tracked on average 395±235 days (range: 46-1089, fixes obtained every 20 min.). All details about capture and movement data are provided in Noonan et al. (2021).

### Data analyses

#### Estimating survival rates

We used the GPS-tracking data to estimate giant anteater survival rates. We considered two groups of resident anteaters, distinguishing those anteaters whose centroid of the home range was far (> 2 km) from paved roads (G_1_, n=20) and near (< 2 km) paved roads (G_2_, n=24). This distance threshold was selected based on Noonan et al. (2021), in which we found that resident giant anteaters with home range centroid distanced > 2 km from a main road had a significantly lower probability of crossing the road, and consequently to be road-killed. For our calculations, we did not consider the dispersal periods but included the data of the two anteaters that settled in new territories (1 month after stopping the dispersal). Previous research showed that there are no significant differences in habitat between roadsides or across roads (Noonan et al., 2021). Estimates of survival were performed using the Kaplan-Meier estimator (Rich et al., 2010) through the R package ‘survival’ (Therneau, 2021).

### Estimating the proportion of the population being road-killed

To estimate the proportion of the population that is road-killed annually, we calculated the real number of roadkills and the number of anteaters inhabiting road vicinity areas.

#### Estimating the real number of roadkills

In Ascensão et al. (2021), we provided an estimate of real roadkill numbers corrected for persistence and detectability biases, using the GENEST framework (Dalthorp et al., 2018). Therein, we used values for the ‘density weighted proportion’ (DWP) i.e., the proportion of total mortality expected to be within the searched corridor area, ranging between 0.80 and 0.95, which seemed reasonable according to our experience in roadkill surveys. Here, we updated these estimates for giant anteater using less-conservative values for DWP. Based on our ongoing movement data collection, we were able to obtain new data on the fate of the anteater carcass location after being hit by a vehicle. Out of the eight confirmed roadkills, five of the collared anteaters were found far (100-600 m) from the road, which corresponds to a DWP of 0.38 (i.e., 38% of the anteater carcasses would be found within the searched corridor area during roadkill surveys). Considering only those road-killed in paved roads, three anteaters out of six still were able to move away from the paved road (up to 100m), which represents a DWP of 0.50 (i.e., half of the carcasses remain within the search corridor area).

This data suggests that a larger proportion of anteaters can move away from the road after colliding with a vehicle. Yet, dirt roads have less traffic (especially trucks) and vehicles travel at a slower speed compared to paved roads. Hence, it is likely that the collisions occurring on dirt roads cause less serious injuries to the animals, and consequently more animals can move away from the collision site. As so, assuming DWP=38 may be overestimating the number of animals moving away from roads after being hit by a vehicle. Hence, we relaxed the DWP parameter and produced estimates of giant anteater roadkill rates using a DWP=0.50. For comparison purposes, we also calculated the roadkill rates when usign DWP=0.80 (Ascensão et al., 2021).

#### Estimating the population abundance

To estimate the population size prone to be affected by roadkill, we needed to estimate the population density and to delimit the road vicinity area i.e., the area in which resident anteaters were more likely to cross the roads. For the population density parameter, we performed a systematic review of the published literature to obtain values for giant anteater population density. We searched the ISI Web of Knowledge and Google Scholar databases, using the search string “ALL FIELDS: (“Giant anteater” OR “Myrmecophaga tridactyla” OR “Tamanduá-bandeira” OR “Tamanduá bandeira”) AND ALL FIELDS: (Density OR Abundance OR Densidade OR Abundância)”, for the timespan 2000-2021. This resulted in 22 scientific articles in ISI Web of Knowledge, and ca. 1750 results in Google Scholar from which we screened the first 10 pages. We reviewed all abstracts and retained only studies that provided some measure of giant anteater density. The reference list of each article was also checked for other relevant publications.

We also included our own estimate using the tracking data. The density estimation was based on the data from one of the study areas (MS040), in an area where an effort was made to capture all resident individuals (Fig.1F). Therein, there were 12 resident giant anteaters, which home ranges overlapped, arranged within a minimum convex polygon of 44.3 km^2^ area. This represents ca. 0.3 individuals per km^2^ (assuming all resident individuals were captured; see Noonan et al. (2021) for details on calculating home ranges).

Finally, we estimated the number of anteaters prone to be affected by roadkill based on density estimates, using a bandwidth of 8 km along roads (4 km for each side) as a conservative distance up to which resident anteaters were likely to cross the roads and be road-killed. This is approximately the longest distance from the road that we have detected animals being tracked that crossed the studied roads.

### Population viability analysis

We used the Vortex model previously described in Desbiez et al. (2020). We ran different sets of simulations, varying the initial population size based on results from previous section (see Results), and the number of animals removed by roadkill per year (based on estimates of real WVC using different DWP values). We modeled a total of 14 scenarios: 3 levels of density values (0.2, 0.3, and 0.4 ind./km^2^), 2 levels of roadkill rates (using DWP of 0.80 and 0.50) and including/not including catastrophes in simulations. We compare the results with the baseline scenario using the minimum population abundance, with and without catastrophes, but without harvesting individuals due to roadkill (Supplementary Material S1). All input parameters are shown in Supplementary Material S2. We used the Vortex version 10.5.5 (Lacy and Pollak, 2021) to generate the PVA models.

## Results

### Survival rates

The yearly survival of giant anteaters inhabiting areas far from the road i.e., with home range centroids distancing >2km from the nearest paved road (G_1_) was estimated to be 0.95 (CI: 0.86-1.00) (Fig.2). For those anteaters inhabiting road vicinity areas (G_2_), the estimated survival decreased to 0.78 (0.62-0.97) (Fig.2). This suggests that the probability of survival of a giant anteater is considerably lower when inhabiting areas close to roads.

**Figure 2.**
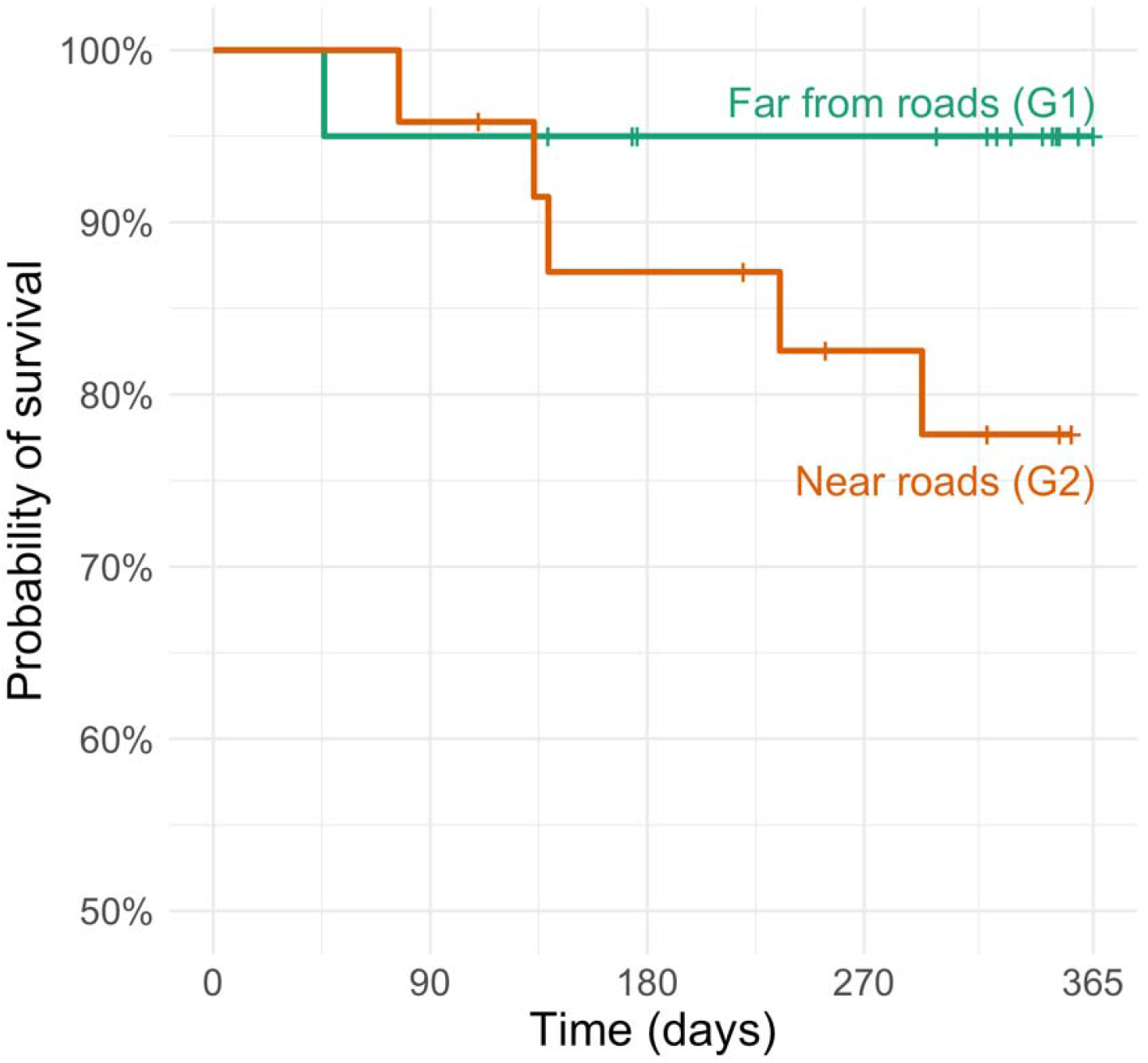
Survival curves for the three groups of giant anteaters tracked within the ‘Anteater and Highways’ research project (www.giantanteater.org). All anteaters (n=42) were residents throughout the tracking period. G_1_ – anteaters living far from roads (n=18), G_2_ – anteaters living near roads (n=24). Survival was estimated using the Kaplan-Meier estimator (Rich et al., 2010).

### Estimating the real number of roadkills

The medians of the estimated real number of collisions involving anteaters when using DWP=0.50 was 1538 (95% CI: 1417-1683), representing ca. 48 (44-52) anteaters per 100 km per year. For comparison, the estimates when using DWP=0.80 were 965 (894-1046), representing a roadkill rate of ca. 30 (28-33) anteaters per 100 km per year, in both cases considerably higher than the roadkill rate obtained from raw values (19 ind./100 km/year).

### Estimating the population abundance

Seven studies reported density values or equivalent (Table S3.1 in Supporting information S3). The density of giant anteater as found in these studies ranged between 0.1 and 2.9 ind./km^2^. However, considered as a reasonable bounding range density of 0.2-0.4 ind./km^2^ (see Supporting information S3). The density of giant anteaters (residents) inhabiting the vicinity of the surveyed roads would therefore range between 160 and 320 individuals per 100 km of roads, when considering densities of 0.2 to 0.4 individuals per km^2^, respectively.

The median of the estimates across studies in savanna areas is 0.3 ind./km^2^, which is the same value we found based on the movement data. Hence, we considered 0.3 ind./km^2^ to be the most probable value of density across the giant anteater distribution range. Combining the roadkill estimates with these abundance estimates, we projected that roadkill is removing annually ca. 20.0% (5-95% CI: 18.5-21.9) of the local population in the vicinity of the road (considering the roadkill estimates using DWP=0.50 and a population density of 0.3 individuals per km^2^) (Fig.3A). The estimates using the range of values of DWP and density varied between ca. 9.4% (8.7-10.2%), when using DWP=0.8 and density=0.4; and 30.0% (27.7-32.9%), when using DWP=0.4 and density=0.2 (Fig.3A).

**Figure 3.**
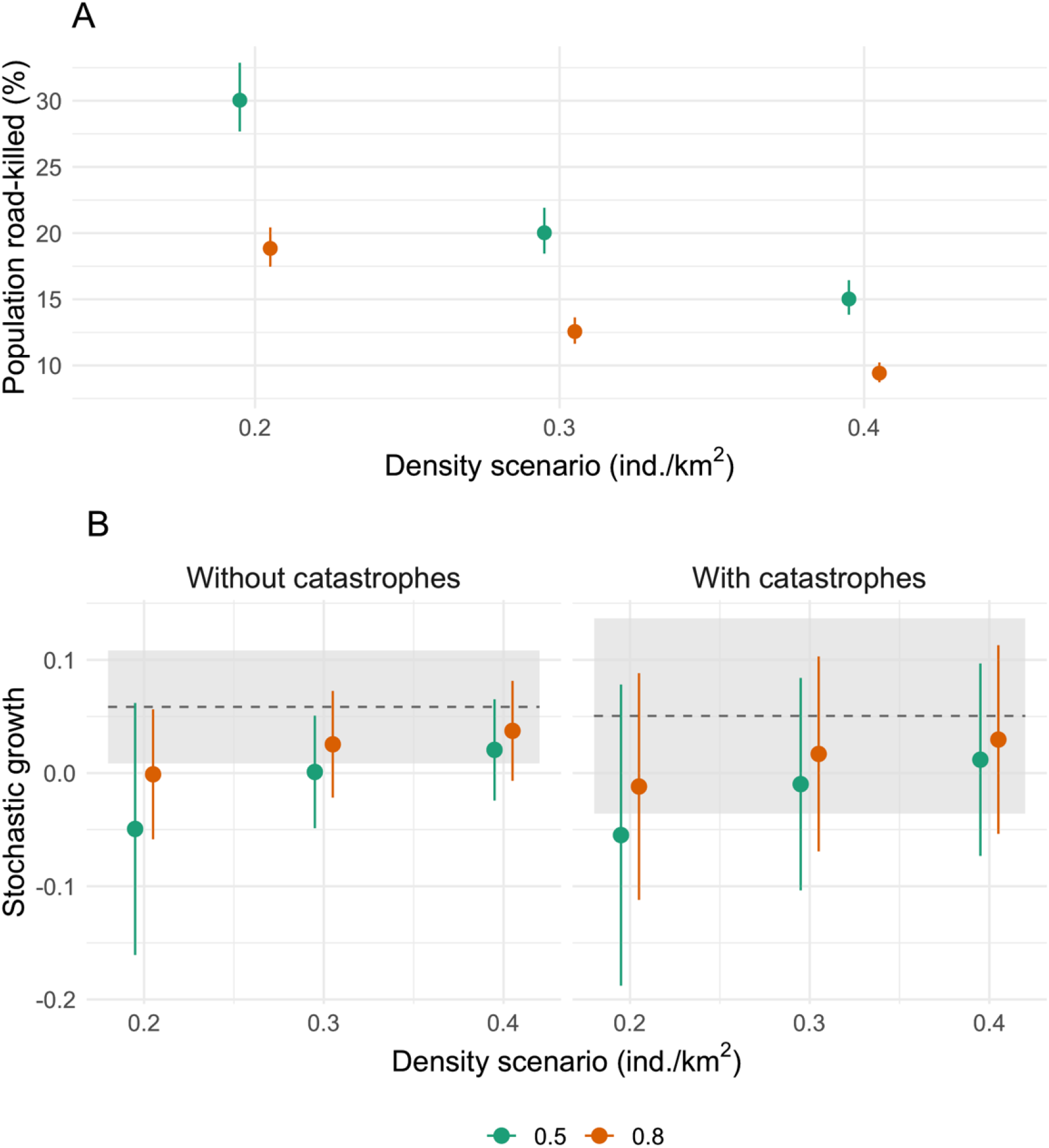
Estimates on the proportion of the population road-killed (A) and of the stochastic growth rate±SD (B) according to different density scenarios (0.2-0.4 ind./km^2^) and value for the parameter DWP (0.5 and 0.8, for calculating total mortality; see text for details). In B) the grey areas and dotted lines represent the stochastic growth rate and SD, respectively, for the baseline scenario (using a density of 0.2 ind./km^2^). In panel A, dots are estimates of median and bars the CI 5-95%. In B), dots are mean values across VORTEX simulations (n=1000) and bars represent SD.

### Population viability analysis

Regarding the results from PVA, the mean±SD stochastic growth rate (*stoch-r*) across the baseline scenario without harvest of individuals from the population by roadkill and in the absence of catastrophes was 5.85±4.98%. When considering the most likely density value of 0.3 individuals per km^2^, and a DWP=0.50, we obtained a mean *stoch-r* of 0.10±4.97% (Fig.3B). The different scenarios of the proportion of individuals road-killed and of the resident population size led to *stoch-r* raging between −4.94±4.98% when density is 0.2 ind./km^2^ and DWP=0.5, and 3.73±4.41% when density is 0.4 ind./km^2^ and DWP=0.8. In all cases, the *stoch-r* when considering roadkill harvesting is considerably lower than the one in the baseline scenario (Fig.3B). When including catastrophic events in simulations, all *stoch-r* values decreased substantially together with higher amplitude in the confidence intervals, denoting a higher vulnerability to local extinctions as well as a higher uncertainty on population trend (Fig.3B). Importantly, all simulations – even without catastrophes – showed a steady reduction in population size, except the simulations using population densities of 0.4 ind./km^2^ and DWP=0.80 (Fig.S4.1 in Supplementary Material S4).

## Discussion

Our results suggest that the survival of resident anteaters is considerably reduced when inhabiting road vicinity areas. Moreover, we estimated that ca. 20% of the population inhabiting road vicinity areas is likely to be extirpated due to vehicle collisions every year. Finally, when modeling the viability of anteater populations nearby paved roads, we obtained a significant decrease in the stochastic growth rate, down to nearly zero (when using density 0.3 ind./km^2^ and DWP=0.5). Therefore, our results clearly support that roads are sink areas, with populations living in road vicinity areas being significantly depleted, requiring the continued recruitment of individuals from other territories (sources).

With the expected expansion and increasing density of the road network (Meijer et al., 2018), the cumulative effect of the source-sink dynamics across the region is likely to be translated into a significantly decrease of animal abundance. Moreover, the habitat of giant anteaters is being rapidly reduced in the Cerrado, namely due to the expansion of soja culture and other cash crops (Green et al., 2019). This means that the source of areas to replenish populations in the vicinity of roads is also decreasing. Such depletion impact may further reduce giant anteater population resilience and ability to withstand or recover from other anthropogenic threats, such as fires, impact of pesticides and agrochemicals, conflict with dogs, persecution by people or disease outbreak (Bertassoni et al., 2019; Garcia et al., 2021; Miranda et al., 2014). Furthermore, although we focused our study on this species, it is reasonable to assume that a similar pattern may be found for other large mammals, known to be highly impacted and vulnerable to roadkill (Fahrig and Rytwinski, 2009; Rytwinski and Fahrig, 2012). As such, roads must be accounted in population and landscape conservation management of current and future road networks, aiming to reduce roadkill throughout the road network.

To the best of our knowledge, the installation of fencing connecting to existing road passages constitutes the single best roadkill mitigation measures (Denneboom et al., 2021; Rytwinski et al., 2016), known to successfully promote the movement of many species, including those vulnerable to roadkill and those posing a risk for humans when involved in wildlife-vehicle collisions (Lesbarrères and Fahrig, 2012). The implementation of road fencing should prioritize sites of higher mortality i.e., hotspots (Spanowicz et al., 2020). However, for low density and broad distribution range species, such as anteaters, detecting hotspots of mortality may not be feasible given the high dispersion of occurrences along roads (Ascensão et al., 2017). An alternative approach is to prioritize places that cross areas of greater connectivity i.e., where animals (are more likely to cross the road in their daily or seasonal displacements (Ascensão et al., 2019a). Such approach has the advantage of requiring few base information, while being unaffected by short-term population dynamics that may shift the location of concentration of roadkill (Santos et al., 2017; Teixeira et al., 2017).

To guarantee its effectiveness, fences must be carefully planned and properly installed, also requiring regular inspection, maintenance, and repair, which will raise the associated costs. Yet, when estimating the mitigation costs, one must consider that large animals represent a serious a threat to humans when involved in collisions with vehicles (Ascensão et al., 2021; Conover, 2019). The fact that a large proportion of anteaters are probably unnoticed in roadkill surveys (this study) also implies that the damage in vehicles and human injuries is also higher. Collisions with large species represent a serious risk for human safety material damage (Ascensão et al., 2021; Conover, 2019). For example, in Mato Grosso do Sul, those collisions involving species with the body size of the giant anteater were estimated to have a material (vehicle) cost of ca. US$ 1150 for cars and US$ 530 for trucks. Using the updated estimates of real roadkill events, the estimated material cost on vehicles of collisions involving giant anteaters ascends to US$ 2,400 per km/year, significantly higher than the previous worst estimate (US$ 985 per km/year; following Ascensão et al., 2021). Hence, investments to reduce the number of collisions, namely proper fencing connecting to existing road passages, are likely to pay off in less than 10 years instead of the previously estimated 16-40 years reported in Ascensão et al. (2021).

We note that there are still some uncertainties in our estimates, namely on the roadkill rates and population abundance. Here, we relaxed the parameter DWP from the GENEST, allowing a higher number of undetected carcasses to be accounted when estimating the total mortality. The proportion of those anteaters being tracked and for which we know that the cause of death was roadkill was considerably high (17%), and it is remarkable that half of the animals hit on paved roads (and all the animals hit on dirt roads) were able to move away from the road. If confirmed, this pattern suggest that the number of road-killed anteaters is largely understimated in roadkill surveys. On the other hand, estimating the population abundance is always challenging to accomplish and there is a high uncertainity on the real density of most species, especially when estimates are made for such large geographic areas. Our bibliographic revision, together with our own estimate on giant anteater density, allowed us to bound the most likely density of this species to 0.2–0.4 ind./km^2^. However, an effort should be made to estimate the density of giant anteaters and other species equally vulnerable to WVC, as only with this information we can provide sound estimates on the impact of roadkill on population persistence (Barrientos et al., 2021).

Additional research should be carried out to quantify the impact of roadkill on the persistence of populations for different species with diverse traits e.g., movement behavior or road avoidance, to ensure that mitigation strategies are effective for a variety of taxonomic groups. Our study serves as an important baseline for large mammals in Brazil. If a substantial part of the road networks is not mitigated, namely in developing countries, the occurrence of iconic species such as the giant anteater will no longer be possible in much of its range.

## Acknowledgements

We would like to thank the donors to the Anteaters & Highways Project especially the Foundation Segre as well as North American and European Zoos listed at (http://www.giantanteater.org/). We would also like to thank the owners of all the ranches that allowed us to monitor animals on their property, in particular Nhuveira, Quatro Irmãos and Santa Lourdes ranches. We are grateful to D.R. Yogui, M. Alves, D. Kluyber and C. Luba for field work, and to Marcello D’Amico and Rafael Barrientos for their valuable comments on an early version of the manuscript. Vortex PVA software (Lacy & Pollak 2021) is provided under a Creative Commons Attribution-No Derivatives International License, courtesy of the Species Conservation Toolkit Initiative (https://scti.tools).

## Statements and Declarations

### Ethics declarations

The authors declare that they have no conflict of interest.

### Data Availability

All data is available upon reasonable request to Arnaud Desbiez.

### Author Contribution

FA and AD contributed equally to the study conception, data analysis and manuscript writing. Both authors read and approved the final manuscript.

