## Supplementary material for "Assessing the impact of roadkill on the persistence of wildlife populations: a case study on the giant anteater"

### Supporting information

#### S1 – Scenarios used in Population Viability Analysis (PVA)

Table S1.1 – Scenarios used in PVA simulations, varying the population density (and corresponding initial population size) and the value of density weighted proportion (DWP) for estimating roadkill rate (i.e., the true number of roadkill events). The proportion of harvesting by roadkill (mean and 5-95% CI) is calculated based on estimates of total mortality and population abundance from empirical data.

| Density (ind./km2) | Population size | Roadkill Scenario | Harvesting due to roadkill (%) |
| --- | --- | --- | --- |
| 0.2 | 160 | Baseline | 0.0 |
| 0.2 | 160 | DWP=0.80 | 18.8 (17.5-20.4) |
| 0.2 | 160 | DWP=0.50 | 30.0 (27.7-32.9) |
| 0.3 | 240 | DWP=0.80 | 12.6 (11.6-13.6) |
| 0.3 | 240 | DWP=0.50 | 20.0 (18.5-21.9) |
| 0.4 | 320 | DWP=0.80 | 9.4 (8.7-10.2) |
| 0.4 | 320 | DWP=0.50 | 15.0 (13.8-16.4) |

#### S2 – VORTEX settings

Table S2.1. *Vortex* parameter inputs for the baseline of the giant anteater (*Myrmecophaga tridactyla*) population model, adapted from Desbiez et.al. (2021).

| **Parameter** | **Value** |
| --- | --- |
| **Species description** | |
| Inbreeding Depression |  |
| Lethal Equivalent | 6.29 |
| % due to recessive lethal | 50 |
| EV Correlation between reproduction and survival | 0.5 |
| **Reproductive System** | |
| Reproductive System | Polygynous |
| Age of 1st offspring females | 2 (limited by a function) |
| Age of 1st offspring males | 3 |
| Maximum lifespan | 15 |
| Maximum # Broods/Year | 1 |
| Maximum # Progeny /Year | 1 |
| Sex Ratio at birth in % Males | 50 |
| Density dependent reproduction | No |
| **Reproductive Rates** |  |
| Percentage of adult female breeding per year | Age 2: 15; Age>2: 60 |
| EV | 5 |
| **Mortality Rates** |  |
| Mortality from age 0 to 1 | 20 |
| SD in 0 to 1 mortality due to EV | 5 |
| Mortality from age 1 to 2 | 8 |
| SD in 1 to 2 mortalities due to EV | 1.6 |
| Annual mortality after age 2 |  |
| SD in mortality after age 2 | 1.6 |
| **Catastrophes** | Yes/No |
| **Mate monopolization** |  |
| % Males in breeding pool | 100 |

#### S3 – Density estimates of giant anteater

The lowest density value (0.1 ind./km^2^) was reported in two studies, one of which in areas with small remnants of native vegetation within a matrix of production forestry (*Pinus* sp.) in an region where anteaters are rarely seen (Braga 2010); and the other was carried out in areas of open savannah adjoining plantations of *Acacia mangium*, which apparently attracted anteaters to their interior given the higher density of anteaters found therein (Kreutz et al., 2012). The lack of remnants seems to be related with the lower density of anteaters found outside plantations (Kreutz et al., 2012). On the other hand, the high density found in Kreutz et al. (2012) (2.9 anteaters/km^2^) had no paralleled in any other study, and therefore is highly unlikely to be frequent across the giant anteater range. In fact, densities as high as 0.4 and higher were only reported in studies carried out in well conserved protected areas (Bertassoni et al., 2021; Bolaño et al., 2015; de Miranda et al., 2006; Desbiez and Medri, 2010).

Table S3.1 – Studies reporting density estimates for giant anteaters.

| **Work** | **Densities (ind./km2)** | **Methods** | **Dominant land cover** |
| --- | --- | --- | --- |
| Bertassoni, A., Bianchi, R., Desbiez, A., in press. Camera trap individual identification of giant anteaters to estimate population size and viability. J. Wildl. Manag. Wildl. Monogr. | 0.4 | Camera-traps |  |
| idem | 0.3 | Camera-traps |  |
| Bolaño, C. R., Cortés, L. M., & Avilán, R. Á. (2015). Densidad Poblacional del oso hormiguero gigante (Myrmecophaga tridactyla) en sistemas ganaderos de pore, casanarE. Revista Biodiversidad Neotropical, 5(1), 64-70. | 0.6 | Linear terrestrial transects | All results |
| idem | 1.0 | Linear terrestrial transects | Natural savanna |
| idem | 0.3 | Linear terrestrial transects | Intervened landscapes |
| Braga, F.G. 2010. Ecologia e comportamento de tamanduá-bandeira Myrmecophaga tridactyla Linnaeus, 1758 no município de Jaguariaíva, Paraná. Tese (Doutorado em Engenharia Florestal). Centro de Ciências Florestais e da Madeira, Universidade Federal do Paraná, Curitiba. 116p. | 0.1 | Linear terrestrial transects |  |
| de Miranda, G. H., Tomas, W. M., Valladares-Padua, C. B., & Rodrigues, F. H. (2006). Giant anteater (Myrmecophaga tridactyla) population Survey in Emas National Park, Brazil--a proposed monitoring program. Endangered Species Update, 23(3), 96-104. | 0.4 | Linear terrestrial transects |  |
| idem | 0.2 | Linear aerial transects |  |
| Desbiez, A. L. J., & Medri, Í. M. (2010). Density and habitat use by giant anteaters (Myrmecophaga tridactyla) and southern tamanduas (Tamandua tetradactyla) in the Pantanal wetland, Brazil. Edentata, 11(1), 4-10. | 0.2 | Linear terrestrial transects |  |
| Kreutz, K. , Fischer, F. , & Linsenmair, K. E. (2012). Timber plantations as favourite habitat for giant anteaters. Mammalia, 76, 137–142. 10.1515/mammalia-2011-0049 | 0.1 | Road counts | In savannah |
| idem | 2.9 | Road counts | In plantations |
| Polisar, J., Scognamillo, D., Maxit, I. E., & Sunquist, M. (2008). Patterns of vertebrate abundance in a tropical mosaic landscape. Studies on Neotropical Fauna and Environment, 43(2), 85-98. | 0.4 | Linear terrestrial transects |  |

#### S4 – Population Viability Modelling – main results

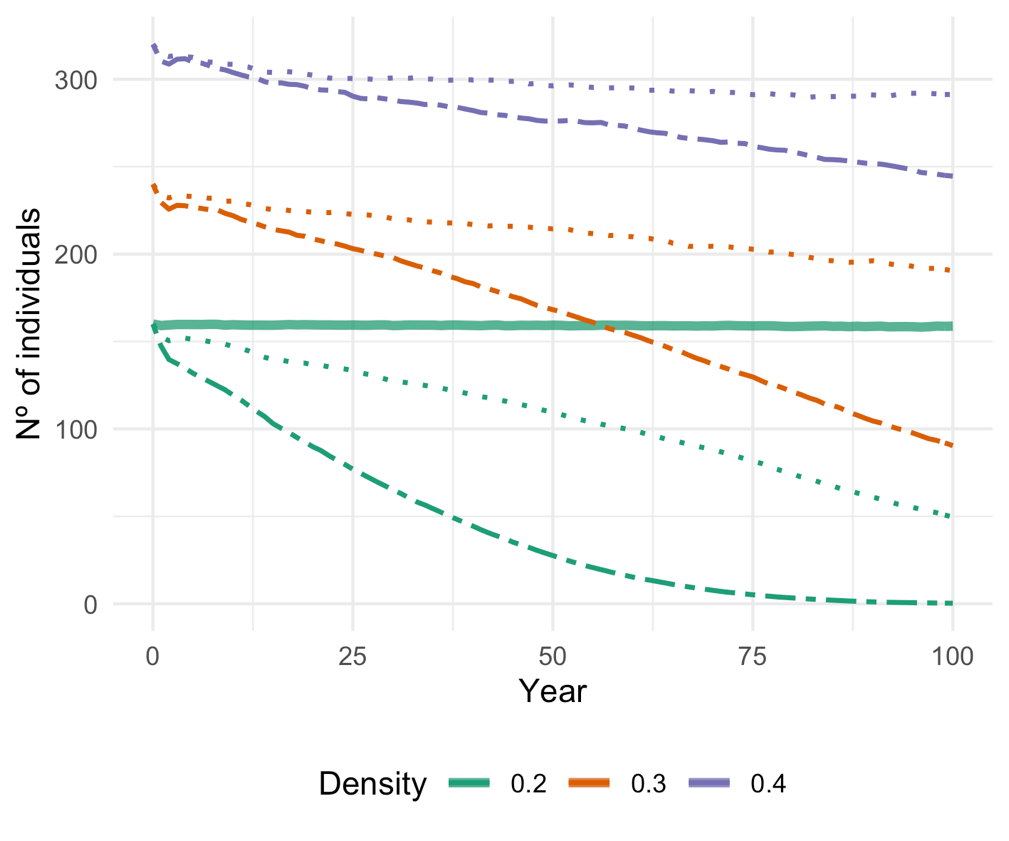

Figure 4.1 – Number of giant anteaters per year, as simulated in the population viability model without catastrophes. Colors denote the population density used (i.e., initial population size), ranging between 0.2 and 0.4 ind./km^2^, and line shape indicate the value of density weighted proportion (DWP) for estimating roadkill rate (i.e., the true number of roadkill events), ranging between 0.5 (dashed lines) and 0.80 (dotted lines). The solid line indicates the baseline scenario, without roadkill harvesting and using a population density of 0.2 ind./km^2^. See main text for details.
